# Ploidy alters root anatomy and shapes the evolution of wheat polyploids

**DOI:** 10.1101/2025.07.11.663263

**Authors:** Jagdeep Singh Sidhu, Harsimardeep S. Gill, Samuel Walker, Harini Rangarajan, Ivan Lopez-Valdivia, Mandeep Singh, Ruairidh J.H. Sawers, Sunish Sehgal, Jonathan P. Lynch

**Affiliations:** Dept. of Plant Science, The Pennsylvania State University; University Park, PA, 16802, USA; Dept. of Agronomy, Horticulture, and Plant Science, South Dakota State University; Brookings, SD, 57007, USA

## Abstract

Polyploidization played a crucial role in crop domestication and modern agriculture. While increased cell size in polyploids is known to enhance plant biomass and vigor, its impact on soil exploration remains poorly understood. Using wheat as a model, we identify a ploidy-induced belowground domestication syndrome, characterized by (a) increased root cortical cell size reducing root respiration, nitrogen content, and phosphorus content; (b) enlarged metaxylem vessels, increasing axial hydraulic conductance; and (c) blunter root tips, limiting penetration ability in compacted soils. Our empirical and *in silico* experiments show that reduced root respiration and reduced cellular nutrient content in wheat polyploids improved nutrient use and acquisition efficiency under suboptimal nitrogen and phosphorus availability. These adaptations would have been advantageous in nutrient-depleted agroecosystems of the Pre-Pottery Neolithic B (PPNB) Fertile Crescent, where continuous cultivation depleted soil fertility over time. Functional-structural modeling indicates that larger cortical cells in wheat polyploids increase vacuolar occupancy, reducing root metabolic costs. Enhanced axial hydraulic conductance may have improved water transport, an advantage in irrigated PPNB agroecosystems. However, polyploids have blunter root tips, which reduces their penetration ability in compacted soils, making them less suited for native soils with greater mechanical impedance. We propose that root anatomical changes driven by ploidy played an important role in adaptations of wheat domesticates to PPNB agriculture.

**One-Sentence Summary:** Polyploidy induced changes in root anatomy may have improved adaptation of wheat to Neolithic agroecosystems.

## Introduction

Polyploidization shaped the evolutionary history of plants and their symbionts (Wendel, 2015; Zhang et al., 2019). Genomic comparisons indicate that all angiosperm species have evolved from one or more episodes of polyploidization, making all flowering species paleo-polyploids (Wood, 2009; Wendel, 2015; Heslop-Harrison et al., 2023). More than 30% of cultivated crops are neo-polyploids (Zhang et al., 2019) including major food crops such as bread wheat (Matsuoka et al., 2011), banana (D’Hont, 2005), *Brassica* sp. (Chalhoub et al., 2014), cotton (Li et al., 2015), sugarcane (D’Hont, 2005), potato (Yang et al., 2017), coffee (Cheng et al., 2017), and many more (Kyriakidou et al., 2018). Given the importance of polyploidy, many aspects of polyploidization have been extensively studied including the underlying polyploid formation mechanisms (Husband, 2004), population genetics (Husband and Schemske, 1997), cytogenetics (McFadden and Sears, 1944), and ecological effects (Levin, 1983) of polyploids. However, the impacts of polyploidy on soil exploration during crop evolution and domestication in Neolithic agroecosystems, particularly in relation to root function, remain largely unknown.

Polyploidization increases DNA copy number, providing advantages such as enhanced genetic diversity, greater vigor, and improved adaptability (Te Beest et al., 2012; Yang et al., 2019). It also increases the probability of novel gene function diversification (Soltis et al., 2014). However, this genomic expansion comes with a potential cost: an increased demand for nucleic acid constituents, particularly phosphorus and nitrogen (William and Lewis, 1985; Neiman et al., 2013; Kang et al., 2015). Nucleic acids (including DNA and RNA) account for up to 40% of cellular phosphorus (Veneklaas et al., 2012) and 5% of cellular nitrogen (Melino et al., 2018; Sidhu et al., 2023). Consequently, polyploids may have greater nutrient requirements than their diploid counterparts (Veneklaas et al., 2012; Anneberg and Segraves, 2023). Under nutrient-limited conditions, increased nutrient demand could place polyploids at a fitness disadvantage, particularly in environments with suboptimal nitrogen and phosphorus availability (Šmarda et al., 2013; Kang et al., 2015; Lynch, 2022). For example, in the freshwater snail *Potamopyrgus antipodarum*, phosphorus limitation causes a two-fold reduction in growth in tetraploids, whereas triploids exhibit no significant change (Neiman et al., 2013). This suggests that the metabolic cost of additional genome copies can be a critical factor influencing polyploid success in nutrient-limited ecosystems.

The hypothesis that plant polyploids have reduced fitness under nutrient-limited conditions appears to contradict their widespread success across diverse soil environments (Wood et al., 2009). This paradox raises key questions about the adaptive and evolutionary mechanisms underlying polyploid fitness, particularly in crop domestication, where higher ploidy levels—such as tetraploid and hexaploid wheat species—were favored (Feldman, 2001; Fuller, 2007; Gegas et al., 2012). In this study, we investigate the adaptive tradeoffs of polyploidy in nutrient-limited environments. A prime example is the Pre-Pottery Neolithic B (PPNB, ∼10,000 Cal-y BP – 7500 Cal-y BP) Fertile Crescent agroecosystems, where continuous land cultivation, evolving irrigation practices, and early agronomic techniques mitigated water deficit stress but gradually depleted soil fertility (Araus et al., 2007, 2014).

Early PPNB (∼10,000 Cal-y BP) farmers in the Fertille Crescent are believed to have settled near river basins (such as Jordon, Euphrates, and Tigris river) and alluvial plains, responding to the region’s arid climate following the Younger Dryas era (∼13,500–11,800 Cal-y BP) as evidenced by archaeological sites such as Tell Abu Hureyra, Dja’de, Cafer Hoyuk and Navali Cori (reviewed in Feldman and Levy, 2023). Isotopic analyses of nitrogen (δ15N) composition in ancient cereal seeds from the Fertile Crescent (Upper Mesopotamia) suggest agriculture emerged (∼9500 Cal-y BP) in favourable environmental conditions including wet and fertile soils but experienced progressive nutrient depletion over the next millennia (Araus et al., 2014). Factors such as continuous cultivation, soil erosion, expansion into less fertile soils, and insufficient manure application likely contributed to this decline. Given these conditions, it is reasonable to assume that successful crop species cultivated in Fertile Crescent evolved adaptive strategies to thrive in irrigated yet nutrient-depleted environments.

Tetraploid wheat (*Triticum turgidum* subsp., 2n = AABB) and hexaploid wheat (*Triticum aestivum,* 2n = AABBDD) are prominent polyploid success stories, both originating in the Fertile Crescent and nearby areas (Eckardt, 2010; Feldman and Levy, 2023). Throughout wheat domestication, farmers selected larger-seeded varieties, including both diploid and polyploid species (Feldman, 2001; Fuller, 2007; Eckardt, 2010; Gegas, 2012; Feldman and Levy, 2023). Einkorn wheat (*Triticum monococcum*, 2n = AA) was the first wheat species to be domesticated in the Fertile Crescent, with the earliest evidence of its cultivation dating around 11,000-10,000 Cal-yBP in upper Mesopotamia near the Karacadağ mountains (Heun et al., 1997). Around the same time (∼10,000 Cal-yBP) emmer wheat species *T. t.* subsp*. dicoccon* was domesticated likely in the north Levant, with the most promising site of origin near the south foothills of the Mt. Hermon (Nave et al., 2019). By the middle PPNB (∼9,800 Cal-Y BP), domesticated emmer wheat had already spread east of the Fertile Cresecent, up to the Caspian belt (Riehl et al., 2013). Thus, cultivated emmer (*Triticum turgidum* subsp. *dicoccon*) emerged as the dominant species in the PPNB, surpassing diploid wheats, likely due to its larger seed size and agronomic advantages (Feldman and Levy, 2023; Gegas et al., 2012). In the South Caspian Sea (likely Mazandran province), cultivated emmer (either dicoccon or parvicoccum) came in contact with *Aegilops tauschii* (2n = DD), and their hydridization gave rise to hexaploid wheat (*Triticum aestivum,* 2n = AABBDD, Zhou et al., 2021). The addition of the D genome to domesticated emmer wheat provided several advantages, including high molecular weight glutenin subunits contributing to superior baking quality, as well as adaptive alleles from Central Asiatic climates that extended the range of bread wheat into the more continental environments of Asia and Europe. While these factors contributed to bread wheat being the most successful wheat species throughout the Holocene and into the modern era, they likely do not represent the full benefits conferred by polyploidy ( Zohary et al., 2012; Smith et al., 2015; Feldman and Levy, 2023). While the aboveground advantages of polyploidy in wheat are well established, the effects of polyploidy on root function and resulting consequences for edaphic adaptation to PPNB agroecosystems are poorly understood.

In this study, we demonstrate that a ploidy-driven increase in wheat cortical cell size led to a belowground domestication syndrome, characterized by larger root cortical cells, wider metaxylem vessels, and blunter root tips. Increased cortical cell size is associated with reduced tissue nutrient content and reduced root respiration, which in turn reduce the metabolic cost of soil exploration. Thereby improving both nutrient capture and nutrient use efficiency. Building on these findings, we propose a novel theory— “Polyploidy is Cheap”—which suggests that polyploidization-driven increases in cell size reduce tissue nutrient demand, making polyploidy an energetically favorable strategy. Additionally, polyploidization leads to larger metaxylem vessels, which increases hydraulic conductance, providing an adaptive advantage in irrigated fields. However, polyploidization also increases root tip bluntness, thereby reducing root penetration in compacted and dry soils, although this effect may not be detrimental in irrigated soils. Local adaptation of the domestication syndrome phenotypes is tested by reconstructing PPNB native and cultivated environments using functional-structural modeling. These results suggest that ploidy-driven root anatomical modifications were key adaptations to the edaphic conditions of early wheat domestication.

## Results

### Increased ploidy is associated with increased nutrient use efficiency

Increased ploidy in wheat is associated with increased cortical cell size (Fig. 1A and B). Tetraploid and hexaploid wheat species had 84% (*P* < 0.0001) and 98% (*P* < 0.0001) larger cortical cells, respectively, compared to diploid species (Fig. 1A and C).

**Figure 1.**
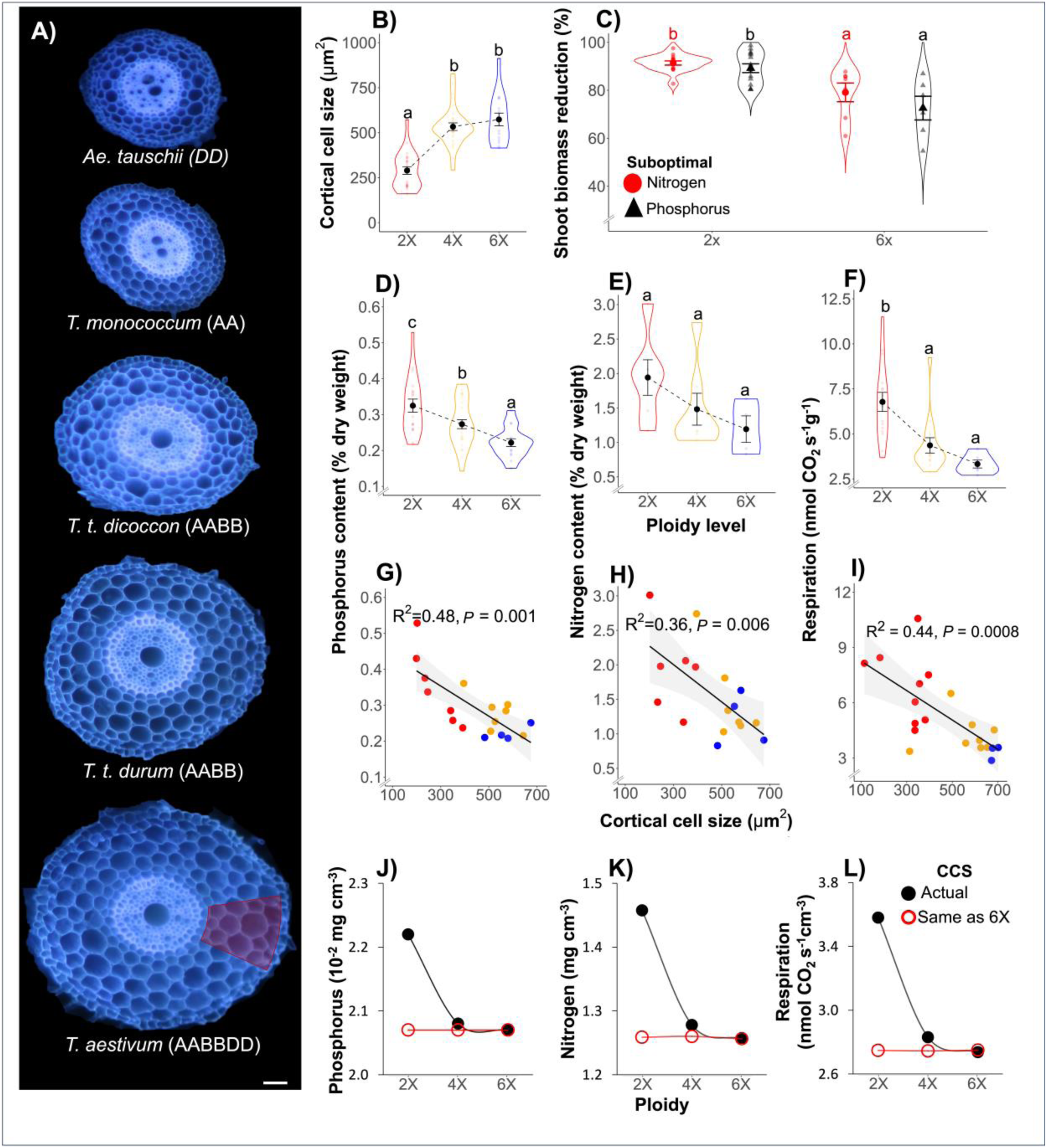
Increased cortical cell size due to polyploidy is associated with greater nutrient use efficiency. Cross-sectional root anatomical images of wheat and its ancestral species with varying ploidy (A), the red shaded section on *T. aestivum* represents cortical cells. Increased ploidy is associated with increased cortical cell size (B). Increased ploidy is linked with reduced sensitivity (represented here as a relative reduction in biomass) to suboptimal nitrogen (red) and phosphorus (black) availability (C). Increased nutrient use efficiency is possibly due to polyploidy driven cortical cell size (CCS) increase (B) which is associated with reduced root tissue phosphorus (D, G), nitrogen content (E, H), and respiration (F, I). *RootSlice* simulation results demonstrate the importance of cortical cell size in this context, as the effect of ploidy on root phosphorus (J), nitrogen (K), and respiration (L) is nullified if all ploidies are simulated with the cortical cell size phenotype of a hexaploid wheat (red), compared to the respective actual phenotypes (black). Each point in panels B–F represents a single biological replicate, with three biological replicates per genotype. In panels G–H, each point represents the average across three biological replicates per genotype. In B-F, the violin plots represent the spread of the data, with black circles indicating means and bars representing standard errors. The letters on top of the violin plots (C-F) denote Tukey’s HSD levels, with groups having different letters being significantly different at p ≤ 0.05. In C, letters denote Kruskal-Wallis test significance. In (G-I) black lines represent best-fitted linear regression lines with grey shading representing the 95% confidence interval around the mean. Scale in panel A = 100 µm.

Under suboptimal nitrogen and phosphorus availability, hexaploid wheat (*T. aestivum*) had 19% (*P* = 0.0005) and 14% (*P* = 0.0007) less reduction in biomass due to suboptimal phosphorus and nitrogen availability, respectively, compared to diploid species (*T. monococcum and Ae. tasuchii,* Fig. 1C).

Increased ploidy was associated with reduced root phosphorus content (Fig. 1D), nitrogen content (Fig. 1E), and root respiration (all measurements per dry weight, Fig. 1F) Relative to diploid species (*Ae. tauschii, T. monococcum*), tetraploids (*T. t.* subsp. *dicoccon*, *T.* t. subsp. *durum*) had 16% (*P* = 0.001) less root phosphorus content, 24% (*P* = 0.33) less root nitrogen content, and 37% (*P* = 0.0001) slower root respiration rate. Compared to diploid species, hexaploid wheat (*T. aestivum*) had 32% less root phosphorus content (*P* < 0.0001), 38% (*P* = 0.13) less root nitrogen content, and 52% (*P* < 0.0001) slower root respiration rate. Differences between tetraploid and hexaploid wheat species were relatively less prominent compared to the diploid species. Nonetheless, hexaploids had 16% (*P* = 0.002) less root phosphorus content (Fig. 1D), 15% (*P* = 0.69) less root nitrogen content (Fig. 1E) and 15% (*P* = 0.23) slower root respiration rate (Fig. 1F) compared to tetraploid wheat species.

Increased ploidy in *Poa* was also associated with reduced root phosphorus content. *Poa annua* (4X) had 16% (*P* = 0.0057) less root phosphorus content than the diploid species *Poa supina* (2X) (Supplementary Fig. S1A). Similarly, *Gossypium hirsutum* (4X) had 25% (*P* = 0.14) less root phosphorus content compared to the diploid species *Gossypium herbaceum* (2X) (Supplementary Fig. S1B).

Increased cortical cell size in wheat across ploidy was associated with reduced nutrient content (Fig. 1G and H) and root respiration (Fig. 1I). A negative linear relationship between CCS and root nutrient content explains 48% (*P* = 0.001) of the variation in root phosphorus content (Fig. 1G), 36% (*P* = 0.006) of the variation in root nitrogen content (Fig. 1H), and 44% (*P* = 0.0008) of the variation in root respiration (Fig. 1I).

To further test the hypothesis that reduced root nutrient content is driven by cortical cell size we simulated the root anatomy of taxa having different ploidy using the functional-structural root anatomical model *RootSlice* (Fig. 1J-L). Using ploidy-specific root anatomical phenotypes, *RootSlice* indicated that increased ploidy is associated with reduced root phosphorus (Fig. 1J), nitrogen content (Fig. 1K), and root respiration (Fig. 1L). Comparing the extreme ploidy levels, the hexaploid phenotype had 7% less phosphorus content, 16% less nitrogen content, and 30% slower root respiration rate than the diploid phenotype. However, when the diploid CCS phenotype was interchanged with the larger CCS phenotype of hexaploid wheat *in silico*, there was no relationship between ploidy and root nutrient content and respiration (Fig. 1J-L).

### Polyploids have greater root axial hydraulic conductance than diploids

Wheat tetraploid and hexaploid species had 22% (*P* < 0.0001) and 18% (*P* = 0.001) larger metaxylem vessel diameter, respectively, compared to diploid species (Fig. 2A and B). The transition from tetraploid to hexaploid did not have any prominent effect on metaxylem vessel diameter (Fig. 2A and B). Estimations of theoretical axial hydraulic conductance from metaxylem vessel diameter show that tetraploids and hexaploids had 81% (*P* < 0.0001) and 78% (*P* < 0.0001), greater axial hydraulic conductance compared to diploid species, respectively, but the addition of the D genome in hexaploid wheat did not result in significant changes in axial hydraulic conductance (Fig. 2C).

**Figure 2.**
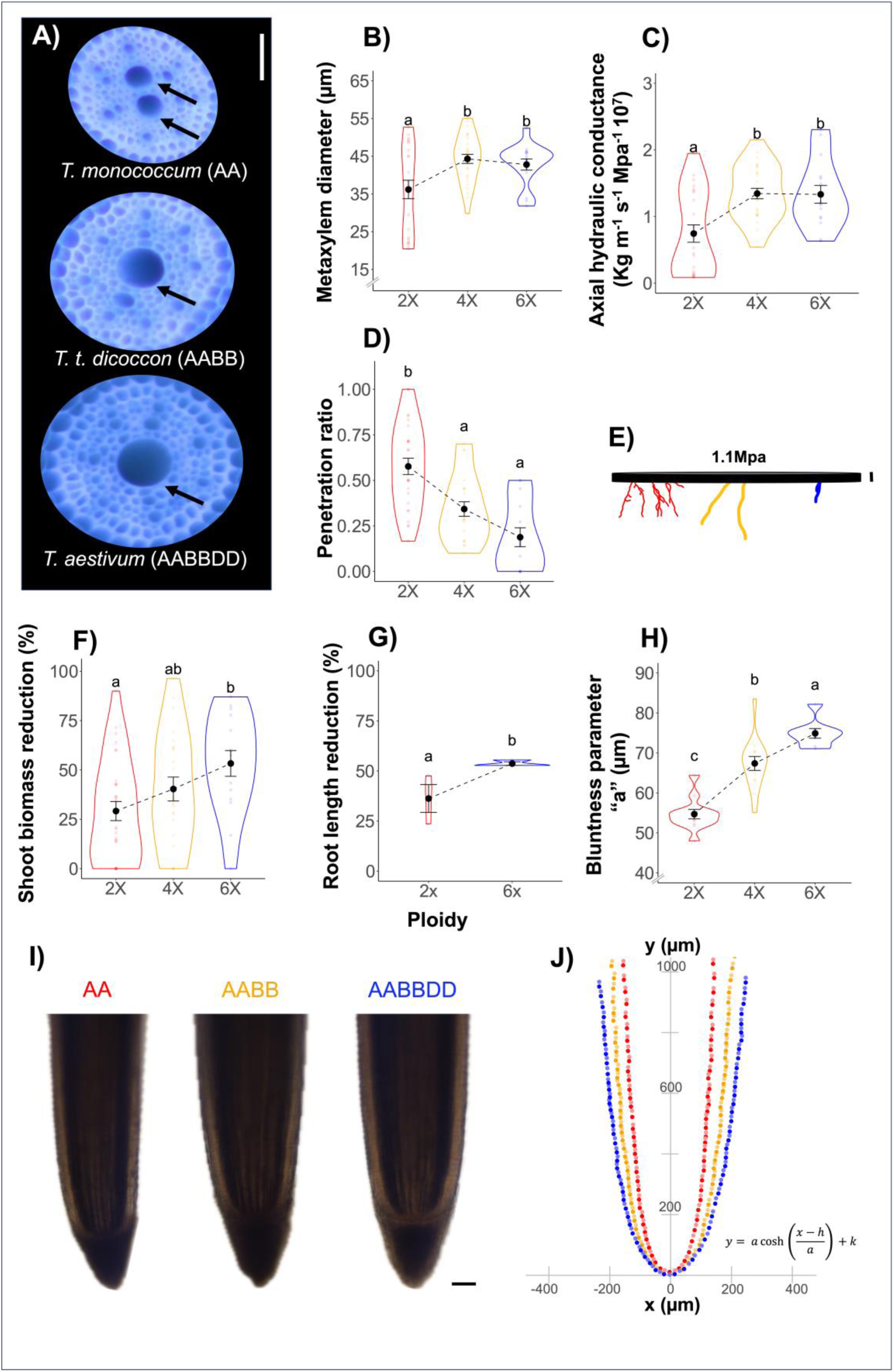
Increased ploidy in wheat is associated with increased axial hydraulic conductance but reduced root penetration ability. Increased ploidy is associated with increased metaxylem vessel diameter (A, B), which increases calculated root axial hydraulic conductance (C). Increased ploidy is associated with reduced root penetration of a hard (1.1MPa mechanical resistance) wax layer (D, E), which impacts shoot biomass production (F), and is also associated with decreased penetration of compacted natural soil (G). Reduced root penetration ability with increased ploidy can be attributed to increased tip bluntness in polyploid wheat (H-J). Representative catenary curves for each ploidy level are shown in J. The parameter “a” defines the bluntness/roundness of a curve with an increase in “a” corresponding to increased bluntness. “h” is horizontal shift (along the x-axis) and “k” is vertical shift (along the y-axis). Arrows in panel A indicate metaxylem vessels. In panels B, C, D, E, G, F and H the letters on top of the violin plots denote Tukey’s HSD levels and in panel G Dunn test levels with groups having different letters being significantly different at p ≤ 0.05. Each colored point in B, C, D, F, G, and H represents a single biological replicate, with three biological replicates per genotype and the violin plots represent the spread of the data, with black circles indicating means and bars representing standard errors. Scales are 50 µm (A), 2 mm (E), and 100 µm (I).

### Increased ploidy is associated with reduced root penetration ability

Greater ploidy was negatively associated with root penetration ability under increased mechanical impedance. Diploid species had 67% (*P* = 0.002) greater root penetration compared to tetraploid species and 216% (*P* < 0.0001) greater root penetration compared to hexaploid species (Fig. 2D and E). Reduced root penetration with increased ploidy was associated with greater reduction in shoot biomass in compacted media, especially comparing hexaploid to diploids (Fig. 2F). Results from the study in which soil impedance was created by a layer of wax:petrolatum were validated by growing diploid (*T. monococcum*) and hexaploid (*T. aestivum*) species in soil that was mechanically compacted to a bulk density of 1.4 g cm^-3^. Confirming the wax: petrolatum results, we observed 48% (*P* = 0.049) greater relative root length reduction due to mechanical impedance in hexaploid species compared to diploid species (Fig. 2G).

The reduced root penetration ability observed in higher ploidy taxa may be associated with increased bluntness of root tips (Fig. 2H and I). The tip shape of all ploidy levels fits well with the catenary curve, with varying “*a*” parameter. The “*a*” parameter quantifies tip bluntness, with lower values indicating sharper tips and higher values corresponding to blunter tips. The average “*a*” value for diploid, tetraploid, and hexaploid species was 54.90, 67.09, and 74.50, respectively (Fig. 2G, I), making tetraploids 22% (*P* < 0.0001), and hexaploids 35% (*P* < 0.0001) blunter than diploids.

### In synthetic polyploids, increased cell size is associated with reduced root nutrient content and root respiration, while blunt root tip shape is associated with reduced penetration ability

The synthetic tetraploid (AADD) had 81% (*P* < 0.0001) greater cortical cell size compared to one diploid parent (*T. monococcum*, AA) and 153% (*P* < 0.0005) greater than its other diploid parent (*A. tauschii*, DD, Fig. 3A and B). Increased cell size in the tetraploid synthetic corresponded with an 87% (*P* < 0.0001) and 90% (*P* < 0.0001) reduction in root phosphorus content compared to AA and DD genome parents, respectively (Fig. 3D). Similarly, the tetraploid synthetic had 53% (*P* = 0.042) and 76% (*P* = 0.0002) slower respiration rate compared to AA and DD genome parents, respectively (Fig. 3F). In contrast, the synthetic hexaploid (AABBDD) had 186% (*P* < 0.0001) larger cells compared to its DD genome parent and 23% (*P* = 0.07) larger cells compared to its tetraploid (*T. t. dicoccoides*, AABB) parent (Fig. 3A and C). As a result, the hexaploid (AABBDD) synthetic had 55% (*P* < 0.0001) less root phosphorus content compared to its diploid (DD) parent but only 10% (*P* = 0.29) less than its tetraploid (AABB) parent (Fig. 3E). Similarly, the synthetic hexaploid (AABBDD) had 51% (*P* < 0.0001) less root respiration compared to its diploid (DD) parent but only 8% (*P* = 0.93) less than its tetraploid (AABB) parent (Fig. 3G).

**Figure 3.**
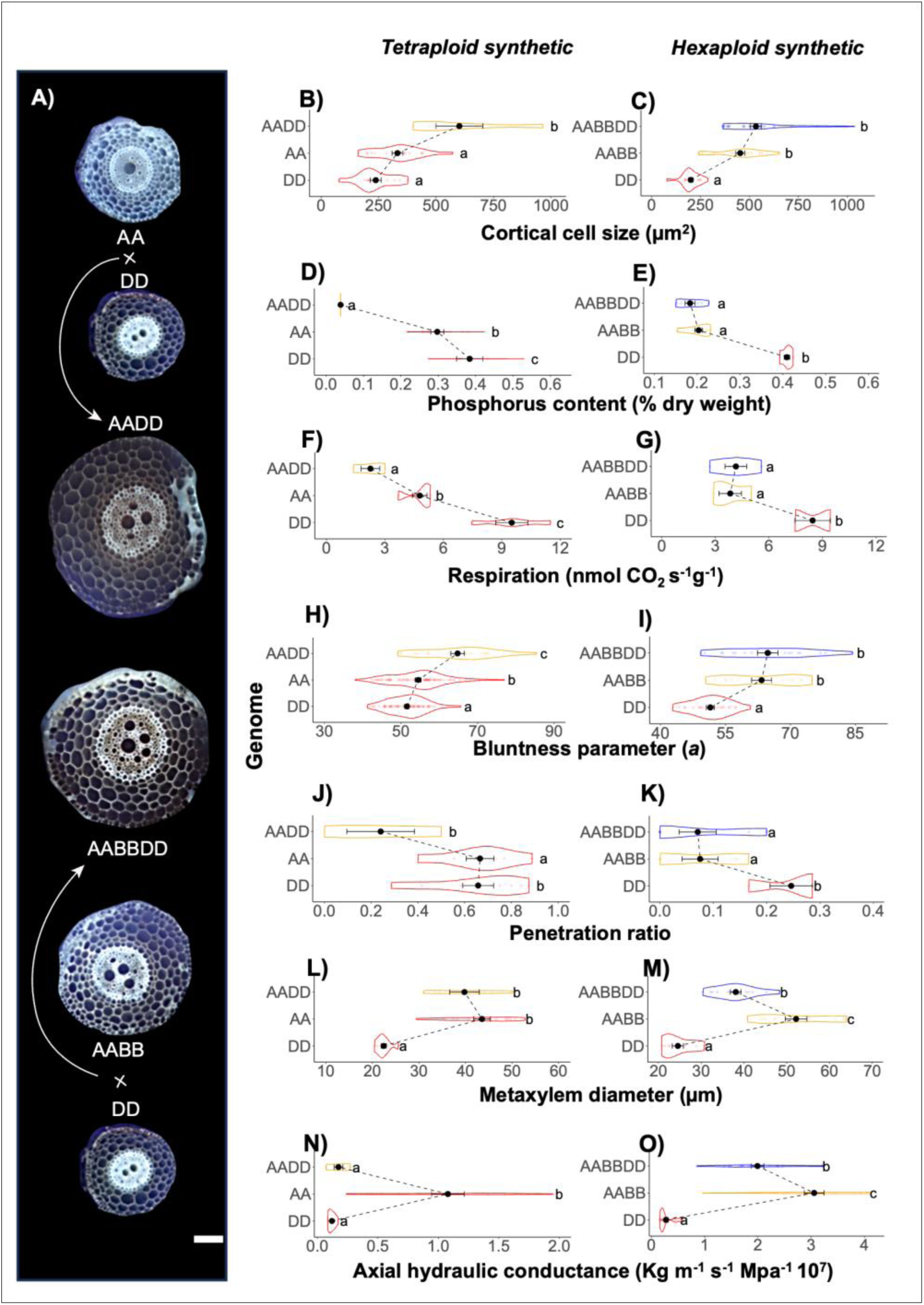
Anatomical and physiological differences in root phenotypes wheat synthetic polyploids. Major changes in root metabolic cost, penetration, and axial conductance in tetraploid synthetics compared to diploid parents but minor in hexaploid synthetics compared to tetraploid parents. Compared to the diploid parents, tetraploid synthetics of wheat have greater cortical cell size (A, B) which is associated with reduced root phosphorus content (D) and root respiration (E), unlike synthetic hexaploids in which increase in cell size is insignificant compared to the tetraploid parents (C), associated with insignificant differences in root phosphorus content (E) and root respiration (F). Synthetic tetraploids have sharper root tip shape (H, represented as bluntness parameter “a” from catenary curve) compared to diploid parents which is likely associated with increased root penetration ability of hard wax layers (J) unlike synthetic hexaploids (I, K). Axial conductance (function of metaxylem vessel diameter and number (L, M)) is less in tetraploid sythetics compared to AA genome parents (N), and less in hexaploid synthetics compared to tetraploid parents (O). The letters on top of the violin plots denote Tukey HSD test levels with groups having different letters being significantly different at p ≤ 0.05. Each colored point in panels B–O represents a single biological replicate, with three biological replicates per genotype. Missing data for any replicate is noted in the accompanying metadata file (available at 10.5281/zenodo.15271839) and the violin plots represent the spread of the data, with black circles indicating means and bars representing standard errors. Scale in C is 100 µm.

The synthetic tetraploid (AADD) had 19% (*P* = 0.012) and 25% (*P* = 0.011) greater root tip bluntness (i.e., values of parameter “a”) compared to its AA and DD genome parents, respectively (Fig. 3H). There was a corresponding 63% (*P* < 0.0001) and 64% (*P* < 0.0001) decrease in the root penetration ratio in the synthetic tetraploid compared to its diploid AA and DD parents, respectively (Fig. 3J). The synthetic hexaploid, in contrast, had 20% (*P* < 0.0001) greater root tip bluntness compared to its diploid (DD) parent but comparable root tip bluntness to its tetraploid (AABB) parent (Fig. 3I). Root tip bluntness was associated with root penetration ability. The synthetic hexaploid had 71% (*P* = 0.029) less root penetration ratio compared to its diploid (DD) parent but comparable root penetration to its tetraploid (AABB) parent (Fig. 3K).

Results for metaxylem vessel diameter did not follow the same trend as for cortical cell size. In synthetic tetraploids, metaxylem vessel diameter was 77% (*P* < 0.0001) larger than one diploid (DD) parent but 8% (*P* = 0.30) smaller compared to its other diploid (AA) parent (Fig. 3L). The synthetic tetraploid exhibited 45% (*P* = 0.91) increased axial conductance compared to its DD genome parent but 83% (*P* < 0.0001) less axial conductance than its AA genome parent (Fig. 3N).

The synthetic hexaploid had 55% (*P* < 0.0001) larger metaxylem vessel diameter (Fig. 3M) and 621% (*P* < 0.0001) greater axial conductance compared to its DD genome parent (Fig. 3O), but 27% (*P* < 0.0001) smaller metaxylem diameter (Fig. 3M) and 36% (*P* = 0.0001) less axial conductance compared to its AABB genome parent (Fig. 3O).

### *OpenSimRoot* modeling of Pre-Pottery Neolithic B Fertile Crescent agriculture

*OpenSimRoot* allowed us to simulate native and cultivated PPNB environments, including soil and atmospheric conditions (Fig. 4A). Simulated plants with polyploid root phenotypes produced greater shoot biomass in cultivated soils compared to plants with diploid root phenotypes, while diploid root phenotypes produced the greatest shoot biomass in native soils (Fig. 4B). Both polyploid phenotypes (4X and 6X) explored a greater soil volume than the diploid phenotype in cultivated environments, whereas the diploid (2X) had better exploration of native soils (Fig. 4C). The hexaploid and tetraploid phenotypes experienced nitrogen and phosphorus stress later and with less severity than the diploid phenotypes in cultivated soils (Fig. 4D). However, in native soils, both polyploid phenotypes experienced nitrogen and water stress earlier and with greater severity than the diploid phenotype (Fig. 4D).

**Figure 4.**
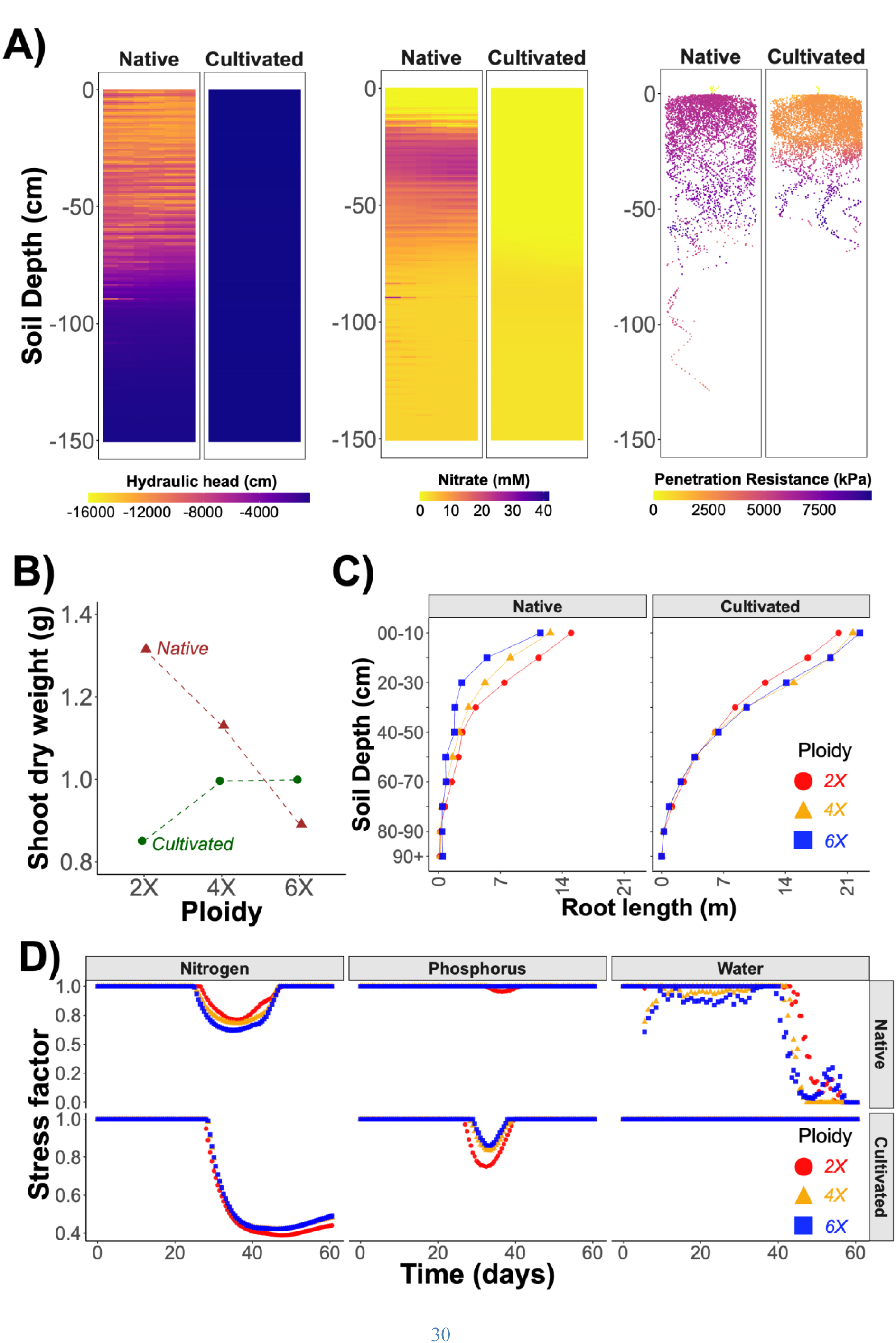
*OpenSimRoot* simulations reveal the adaptive advantages of polyploid wheat root phenotypes in cultivated Pre-Pottery Neolithic B (PPNB) environments (∼7,500 Cal-y BP) and diploid wheat root phenotypes under native (*i.e.*, uncultivated) PPNB environments (∼10,000 Cal-y BP). Root architecture and soil conditions were simulated using the functional–structural model *OpenSimRoot v2* with parameters given in tables S3 and S4. Distribution profile of water, nitrate, and root penetration resistance in PPNB native and cultivated soils (A). Shoot dry weight at 60 days (B). Root length distribution by depth at 60 days (C), and stress factor (1 is for non-stressed plants) for nitrogen, phosphorus, and water over time of different ploidy species of wheat under native and cultivated enviroments. The input files required to reproduce these simulations are available in the following Zenodo repository 10.5281/zenodo.15271839.

## Discussion

In this study, we demonstrate that polyploidization is associated with root anatomical phenotypes that may have played an adaptive role in plant evolution and crop domestication. By utilizing the wheat domestication model, we provide evidence that 1) polyploidy increases root cortical cell size, which is associated with reduced root tissue nutrient content and greater nitrogen, phosphorus, and carbon use efficiency; 2) polyploidy is linked to an increase in root metaxylem vessel diameter and greater axial hydraulic conductance; and 3) increased ploidy is associated with blunter root tips and reduced root penetration of hard soil. Based on these observations, we propose a belowground domestication syndrome for wheat that explains how ploidy-driven root anatomical changes could have played an adaptive role in wheat domestication. Additionally, we discuss the implications of the ploidy-associated reduction in root nutrient content and increased nutrient efficiency in polyploids.

The question of whether polyploidization increases cellular nutrient demand has been debated extensively in both animal and plant literature, with several theories proposing ‘polyploidy is expensive’ (William and Lewis, 1985; Cavalier-Smith, 2005; Šmarda et al., 2013; Neiman et al., 2013; Guignard et al., 2016). For example, it has long been thought that the increased nuclear DNA content resulting from either increased genome size or whole genome duplication would be disadvantageous to an organism due to the increased nutrient requirement of additional DNA (Neiman et al., 2013; Kang et al., 2015; Anneberg and Segraves, 2023). Another theory suggests polyploids would have a greater demand for nutrients not because of extra DNA but because of the development and maintenance cost of larger cells and cytoplasmic machinery (Šmarda et al., 2013). Although these theories are applicable to some plant and animal species (Neiman et al., 2013; Kang et al., 2015; Guignard et al., 2016), they fail to explain the abundance of polyploid plant species, particularly in nutrient-limited ecosystems such as phosphorus-limited arctic regions dominated by polyploids (Brochmann et al., 2004) and nitrogen-limited Neolithic Near East agroecosystems where wheat polyploids evolved (Feldman and Levy, 2023). Furthermore, most studies have overlooked the impact of ploidy on root phenotypes and tissue nutrient demand, despite roots being a significant sink of these scarce resources and the obvious fact that roots are responsible for the capture of soil resources (Nielsen et al., 1994; Nielsen et al., 1998; Lambers et al., 2002; Lynch, 2015; Lynch 2019; Lynch et al., 2021). In this study, we propose a counterintuitive theory, which we support with substantial evidence – ‘*polyploidy is cheap*’ -stating that increased cell size associated with polyploidy reduces both the nutrient and respiratory cost of root tissue, leading to more efficient soil exploration, greater capture of soil resources, and greater efficiency of nutrient use in root tissue.

We show that increased ploidy in wheat is associated with greater phosphorus and nitrogen acquisition and use efficiency (Fig. 1B and 4B) due to reduced root phosphorus and nitrogen content and slower root respiration (Fig. 1D, E, and F). Both empirical and *in silico* results demonstrate that larger root cortical cell size in polyploids reduces root phosphorus and nitrogen content (Fig. 1G, H, I, J, K, and L). These findings contrast with previous ‘polyploidy is expensive’ theories, which treated genome size and ploidy similarly and neglected the role of vacuoles in reducing cellular nutrient demand. Vacuolar compartments are metabolically cheap compared to the cytosol and nonvacuolar organelles (Lee et al., 1990; Dünser et al., 2022; Sidhu et al., 2023; Sidhu and Lynch, 2024) including the nucleus, the endomembrane system, mitochondria, plastids, and in photosynthetic cells, chloroplasts, which collectively are the major sink of phosphorus and nitrogen in cells, and the major respiratory cost (Melino et al., 2018; Veneklaas et al., 2012). Greater vacuolar occupancy in maize root cortical cells is associated with reduced root phosphorus content, nitrogen content, and respiration (Chimungu et al., 2014; Lopez-Valdivia et al., 2023; Sidhu and Lynch, 2024). The ploidy-driven increase in cell size employs the aforementioned mechanism of increased vacuolar: cytoplasmic ratio per cortical volume, which ultimately reduces root tissue nutrient content.

Reduced root nutrient and carbon demand due to increased cell size improves the metabolic efficiency of soil exploration, resulting in greater resource capture under drought and low nitrogen in maize (Chimungu et al., 2014; Lopez-Valdivia et al., 2023) and under increased soil penetration resistance in wheat (Colombi et al., 2019). This improved resource acquisition increases photosynthesis, which supports greater shoot growth, enabling further root development—establishing a positive feedback loop that reinforces whole-plant performance in nutrient- and water-limited environments (Chimungu et al., 2014; Lopez-Valdivia et al., 2023; reviewed in Lynch et al., 2021). Our results suggest that wheat polyploids likely benefit from the same mechanism of enhanced soil exploration efficiency, enabling them to outperform diploid wheat under phosphorus and nitrogen limitation (Fig. 1C and 4B). However, these findings warrant further validation in field experiments involving a broader range of diploid and polyploid wheat species.

Physiological comparison of wheat taxa with varying ploidy is potentially confounded by the distinct evolutionary trajectories of these taxa. To overcome this issue we employed two strategies: 1) we evaluated synthetic wheat polyploids recently generated from diploid and tetraploid species, and 2) we used functional-structural modeling to isolate the effects of cell size from other phenotypic differences among wheat taxa.

Analysis of synthetic tetraploid and hexaploid wheat supports the theory that ploidy-driven increases in cortical cell size reduce root nutrient content and metabolic cost (Fig. 3). We observed a significant decrease in root phosphorus content and respiration in synthetic tetraploids, which gained increased cortical cell size upon neopolyploidization (Fig. 4A, D, and F). In contrast, neo-hexaploidization does not result in a significant further increase in cortical cell size compared to tetraploid parents, and consequently does not show decreased root phosphorus content and respiration compared to the tetraploid parents (Fig. 4B, E, and G). It is worth noting that the majority of wheat domestication occurred in tetraploid wheat, and only domesticated tetraploids gave rise to hexaploid bread wheat. Thus, the major increase in cell size was fixed shortly after the tetraploidization event. A similar trend of prominent cell enlargement following tetraploidization, but more limited increases with hexaploidization and octoploidization, has been reported in synthetic *Arabidopsis* polyploids (Tsukaya, 2008). While it is possible that natural and/or artificial selection could have further selected for increased cortical cell size in wheat polyploids (Fig. 1C), our results suggest that the initial increase in cell size is due to polyploidization (Fig. 3A). The use of synthetic polyploids allowed us to control for selection effects, providing strong evidence that polyploidization itself is the primary driver of changes in root cortical cell size and related phenotypes.

To further isolate the effects of ploidy-induced cortical cell size increase on root nutrient content and metabolic cost from other phenotypic differences among wheat taxa, we used a functional-structural modeling approach via *RootSlice*. *RootSlice* is a heuristic model that permits precise functional-structural simulation of root anatomical phenotypes at from root segment to subcellular scales (Sidhu et al., 2023). *RootSlice* has been successfully used to explore hypotheses, test the adequacy of conceptual models, and isolate specific phenotypic effect values (Sidhu and Lynch, 2024; Sidhu et al., 2024). Unlike empirical comparisons, *RootSlice* allows precise manipulation of individual anatomical phenotypes—such as cortical cell size—while holding other variables constant. This enables assessment of how specific anatomical phenotypes affect tissue nutrient content and respiration, independent of noise from the environment, the broader phenotype, and genetic background. *RootSlice* simulations confirmed that increased cortical cell size in polyploids leads to a greater proportion of vacuolar compartment per unit cortical volume and a corresponding reduction in metabolically active cytoplasmic volume (Fig. 1J, K and L). This results in less root phosphorus and nitrogen content, and reduced respiration rates. Furthermore, when all ploidy phenotypes were simulated using the hexaploid cortical cell size phenotype while holding the remainder of the phenotypes to be ploidy specific, the differences in root nutrient content and respiration among ploidy levels disappeared. These results indicate that increased cortical cell size alone can account for the majority of the decrease in root nitrogen, root phosphorus, and respiration in polyploids, providing evidence that the effects of polyploidy on root metabolism are mediated primarily through changes in cell size rather than other ploidy-linked factors. More broadly, our use of *RootSlice* demonstrates the power of functional-structural modeling to mechanistically dissect the impacts of anatomical phenotypes on root function and whole-plant metabolic economy—an approach that complements and strengthens empirical studies.

The phenomenon of polyploidization-associated reductions in root metabolic cost is likely applicable to other plant species, particularly monocots. In a comparative analysis of *Poa* species—namely, *Poa annua* (4X) and *Poa supina* (2X)—we observed similar results, with the tetraploid species exhibiting lower root phosphorus content than the diploid species (Supplementary fig. S1A). In contrast, the tetraploid *Gossypium hirsutum* (AABB) did not show statistically significant differences in root phosphorus content compared to its diploid counterpart, *Gossypium herbaceum* (AA) (Supplementary fig. S1B). This discrepancy is likely due to secondary growth in dicots, which destroys the root cortex in the mature root zone (Strock et al., 2018), thus reducing the benefits of increased cortical cell size. Further confirmation of these results will require evaluation of a larger sample of taxa within each ploidy level. Still, polyploidy (whether endopolyploidy, autopolyploidy, or allopolyploidy) is associated with increased cell size in both monocots and dicots, including model species such as *Arabidopsis* (Tsukaya, 2008), suggesting that the “polyploidy is cheap” hypothesis may apply across plant taxa. The energetic and nutritional advantages of increased vacuolar occupancy and reduced cytoplasmic volume may be especially relevant for early terrestrial plants, including ferns, bryophytes, and lycophytes, which often inhabit nutrient-poor environments (Brundrett, 2002). Further exploration across eukaryotic lineages may reveal that polyploidy and cell size are not found under just low-resource environments, but may be favored evolutionary strategies for reducing metabolic cost and increasing ecological fitness.

Additionally, our theory of ‘*polyploidy is cheap*’ can also explain why ploidy in some animals increases nutrient demand (Neiman et al., 2013), as animal cells do not possess energy-efficient vacuoles. Unlike plant cells, animal cell expansion must rely on metabolic processes that consume greater amounts of energy and nutrients, including the synthesis of proteins and other cytoplasmic machinery that require significant amounts of phosphorus, nitrogen, and energy (Björklund, 2019). Therefore, the increased DNA content associated with ploidy in animals may lead to a greater demand for nutrients to support these processes (Neiman et al., 2013). However, other energy-efficient ways, such as decreased total cell surface area, might be at play, that can reduce tissue metabolic cost in animal polyploids, for example, in *Xenopus* (Cadart et al., 2023).

Increased ploidy in wheat also results in increased metaxylem vessel diameter (Fig. 2B), which is predicted to increase root axial hydraulic conductance (Fig. 2A). Our expected axial hydraulic conductance values are consistent with empirical evidence (Zhao et al., 2005) supporting that increased ploidy in wheat leads to increased axial hydraulic conductance at the scale of root axes and root systems. While increased axial hydraulic conductance can provide functional advantages by increasing water transport under optimal water availability, it can be maladaptive under water deficit due to the associated risk of embolism and increased water loss (Richards and Passioura, 1989; Hendel et al., 2021; Strock et al., 2021). Therefore, increased ploidy in wheat provides a favorable phenotype characterized by larger xylem vessel diameter which could be advantageous in environments with ample water supply but potentially disadvantageous under drought. Archaeological evidence suggests that the domestication of polyploids occurred close to riverbeds and later in irrigated fields of the Fertile Crescent (Araus et al., 2007; Aguilera et al., 2008; Araus et al., 2014; Feldman and Levy, 2023), which would have given an advantage to polyploids over diploids. However, increased root metaxylem vessel diameter in wheat polyploids is not solely due to ploidy, as synthetic tetra and hexaploids don’t have larger metaxylem diameter compared to all their parents (Fig. 3N and O). This indicates that greater axial conductance in tetraploid and hexaploid wheat (Fig. 2A, Zhao et al., 2005) is also due to natural or artificial selection during the domestication of wheat polyploids. The variation in xylem vessel diameter within hexaploid wheat germplasm (Fig. 2A, Hendel et al., 2021) is encouraging as it provides a basis for selecting a smaller root xylem vessel diameter, a phenotype that is useful under terminal drought stress (Richards and Passioura, 1989; Hendel et al., 2021; Strock et al., 2021). It is important to note, however, that axial conductance is not always a rate-limiting step in water transport; rather, soil-root interface resistance and radial conductance through cortical tissue often dominate whole-plant hydraulic resistance (Tyree and Zimmermann, 2002; Steudle, 2000). Nevertheless, smaller metaxylem vessel diameter can be advantageous under drought stress (Richards and Passioura, 1989; Hendel et al., 2021; Strock et al., 2021)

Increased ploidy in wheat is associated with reduced root penetration ability in media with increased mechanical impedance (Fig. 2D, E, and H). While the wax:petrolatum layer assay is a useful high-throughput platform to study root penetration in response to soil mechanical impedance (Long-Xi et al., 1995), in this study we confirmed results with a wax:petrolatum layer by growing plants in natural soil with greater bulk density (1.4 g cm^-3^). We found that diploid root growth is less sensitive to increased mechanical impedance compared to hexaploid taxa (Fig. 2F). We propose that reduced penetration ability in polyploids can be partially explained by the increase in root tip bluntness (Fig. 2G and I). Increased root tip bluntness in wheat increases penetration stress and reduces root elongation in hard soils (Colombi et al., 2017). Additionally, indirect evidence involving soil penetration probes and 3D printed root probes shows that sharper tip shapes experience less mechanical resistance (Ruiz et al., 2016; Mishra et al., 2018). Therefore, it is reasonable to propose that increased root tip bluntness in wheat polyploids contributes to reduced root penetration ability. Increased root tip bluntness observed in wheat polyploids is primarily due to an increase in cortical cell diameter, which leads to an overall increase in root diameter and constrains root tip morphology, resulting in a blunter shape compared to diploids. As with metaxylem vessel diameter and cortical cell size, we observe significant variation in root tip shape (Fig. 2G) and penetration ability (Fig. 2D) within hexaploid wheat germplasm, which can be exploited to improve the root penetration ability of wheat without altering the ploidy level.

By integrating the ploidy-induced changes in root anatomy we propose a belowground domestication syndrome that includes increased cortical cell size (increasing nutrient use and acquisition efficiency), increased root metaxylem vessel diameter (increasing axial hydraulic conductance), and increased root tip bluntness (reducing root penetration of hard soils). The increased nutrient efficiency, increased axial hydraulic conductance, and reduced root penetration ability observed in wheat polyploids are phenotypes that may be well-suited for PPNB (10,000-7500 Cal-y BP) agroecosystems of the Fertile Crescent, in which wheat was domesticated. Early PPNB (∼9500 Cal-y BP) farmers in the Fertile Crescent (especially in upper Mesopotamia) are believed to have settled near river basins and alluvial plains, responding to the region’s arid climate following the Younger Dryas era (∼13,500–11,800 Cal-y BP) as evidenced by archaeological sites such as Tell Abu Hureyra, Dja’de, Cafer Hoyuk and Navali Cori (reviewed in Feldman and Levy, 2023). Indirect evidence based on carbon (δ^13^C) isotope composition of archaeological charred kernels suggests that during the early the Pre-Pottery Neolithic period (∼10,000 to 7,500 Cal-y BP) Fertile crescent crops and associated weed species enjoyed greater soil water availability which gradually declined over time (Araus et al., 2007; Araus et al., 2014). Analysis of the nitrogen (δ^15^N) isotope composition of these samples indicates that the agrarian soils of Mesopotamia were initially fertile (∼10,000 Cal-y BP) but gradually lost fertility over time, presumably due to factors including continuous cultivation, soil degradation and erosion, expansion of agriculture to less fertile soils, and nonexistent or inadequate manure application (Araus et al., 2007; Araus et al., 2014). Numerous other modern and prehistoric examples illustrate how continuous farming, in the absence of sufficient nutrient replenishment, has driven long-term nutrient depletion and progressive alterations to soil structure through mechanical disturbance and soil degradation (Henao and Baanante, 1999; Redman, 1999; Tilman et al., 2002; Kirch and Sharp, 2005; Hartshorn et al., 2006). For instance, approximately 500 years of taro cultivation in Hawai‘i (beginning around 1600 C.E.) resulted in the loss of 42% of the phosphorus from ash horizons that had remained undisturbed by humans for over 17,000 years (Hartshorn et al., 2006).

Irrigation reduces mechanical impedance to root growth (Whitmore and Whalley; 2009; Ruiz et al., 2016; Lynch, 2022) but leads to soil erosion and nutrient leaching, especially nitrate-nitrogen (Graham et al., 2022). Therefore, based on water availability and the alluvial nature of the Fertile Crescent (especially upper fertile Mesopotamia) agricultural soils during the early PPNB period, we can infer that early wheat domesticates did not face substantial mechanical impedance. A recent *in silico* study showed that the advent of irrigation coincident with crop domestication may have changed the fitness landscape of root responses to soil mechanical impedance (Rangarajan and Lynch, 2024). A recent study of root evolution during maize domestication found that anthropogenic changes in soil conditions were associated with root architectural and anatomical adaptations (López-Valdivia et al., 2024). Increasing atmospheric CO₂ concentrations during the early Holocene played an important role in the emergence of reduced nodal root number and multiseriate cortical sclerenchyma, promoting deeper rooting. The introduction of irrigation by ∼6,000 Cal-y BP shifted nitrogen availability from the topsoil to subsoil, further enhancing the functional value of these phenotypes. These results from Neolithic maize domestication environments parallel our observations in wheat, where ploidy-driven increases in cortical cell size may have conferred advantages under similar conditions of reduced mechanical impedance and declining soil fertility during early domestication in the Fertile Crescent.

Using *OpenSimRoot v2* we simulated native and cultivated PPNB ecosystems, setting soil and environmental parameters to mimic archaeological sites from the upper Fertile Crescent, such as Tell Abu Hureyra (Fig. 4A, Supplementary Table S3 and S4). Modeling results confirm that diploid root phenotypes, characterized by better soil penetration ability and reduced axial conductance, are better suited for exploring the dry, hard soils of native environments and produce greater shoot biomass under these conditions (Fig. 4B–D). In contrast, polyploid root phenotypes, which exhibit reduced root construction and maintenance costs and greater axial conductance, are better adapted to cultivated soils with greater water availability and less fertility (Fig. 4B–D). The reduced root penetration ability associated with increased ploidy is less disadvantageous in cultivated alluvial soils, where soil mechanical impedance is substantially less than in native soils (Fig. 4A).

Its worth noting that with *OpenSimRoot v2* simulations, we are strictly testing the value of root phenotypes associated with different ploidy levels (Supplementary Table S2), while controlling for any differences in phenology and shoot growth by using the same base shoot model across all scenarios. The model explicitly incorporates key biophysical processes relevant to PPNB Fertile Crescent conditions, including soil mechanical impedance as a function of soil depth, texture, bulk density and moisture, water and nutrient dynamics, and photosynthetic responses to atmospheric CO₂ levels (270 ppm in the PPNB) and the dynamics of plant and root growth in response to water and nutrient availability (Strock et al., 2022; Rangarajan and Lynch, 2024; López-Valdivia et al., 2024). *OpenSimRoot v2* is a heuristic rather than a predictive model, and in the present context provides quantitative support for the logic model and hypothesis that root anatomical changes associated with polyploidy were positive adaptations to the PPNB agroecosystems in which wheat domestication occurred. Therefore, by reconstructing PPNB agroecosystems where direct measurements are impossible, *OpenSimRoot v2* allowed us to test the utility of PPNB root phenotypes under realistic environmental and management conditions, a task that would be very challenging to test empirically.

Although the prevalent theory is that wheat polyploids may have been selected for their increased seed size (Fuller, 2007), we propose that root anatomical changes induced by polyploidization also provided adaptiveness during the Neolithic period. It is likely that Neolithic farmers directly selected for larger seeds—the most direct and agronomically valuable phenotype—which polyploids naturally possess. However, larger seed size may have come as part of an integrated anatomical package that includes larger root cortical cells and leaf mesophyll cells (Wilson et al., 2021). It is also reasonable that early farmers, knowingly or unknowingly, selected plants that performed better under the nutrient-depleted yet wetter soils typical of Neolithic agriculture (Araus et al., 2014), favoring genotypes with increased cortical cell size that confer improved nitrogen and phosphorus acquisition and use efficiency. As a consequence, larger seed size may have been indirectly selected. Regardless of whether seed size or root anatomy was the primary target of selection, ploidy associated anatomical changes -including increased cortical cell size (reducing root metabolic cost), larger mesophyll cells (enhancing photosynthetic capacity), larger endosperm cell size (increasing seed size), and larger metaxylem vessels (enhancing axial water transport)—constitutes a favorable integrated phenotype for PPNB agroecosystems. These coordinated changes likely contributed to improved yield and whole-plant performance. Importantly, such a complex phenotype could be rapidly achieved as pleiotropic results from polyploidy, offering a compelling explanation for their dominance in modern wheat cultivation.

In conclusion, this study highlights the functional and adaptive significance of polyploidization-induced root anatomical changes, especially during crop domestication. We show that increased cortical cell size in wheat polyploids enhances phosphorus and nitrogen acquisition and carbon use efficiency; enlarged metaxylem vessel diameter is associated with greater axial hydraulic conductance; and greater root tip bluntness— constrained by cortical cell expansion—reduces root penetration in compacted soils. By integrating these phenotypes we define a belowground domestication syndrome for wheat that likely provided competitive advantages to wheat polyploids under the irrigated but nutrient-depleted agroecosystems of PPNB and beyond. Ploidy-dependent changes in root anatomy also offer broader insights into the ecological and evolutionary roles of polyploidy in shaping plant adaptation to resource-limited environments, including drought and low fertility. Moreover, the significant anatomical variation observed within major crops including wheat, maize, and rice germplasm presents an opportunity to breed cultivars with improved resource use efficiency by selecting for larger cortical cells, smaller metaxylem vessels, and sharper root tips—phenotypes that can optimize root function without altering ploidy.

## Materials and Methods

### Root anatomy and tissue nutrient content

For this experiment, we tested four genotypes for each of six species including two diploids (*Triticum monococcum*, *Aegilops tauschii*), two tetraploids (*Triticum turgidum* subsp. *dicoccoides*, *Triticum turgidum* subsp. *dicoccon, Triticum turgidum* subsp. durum) and a hexaploid (*Triticum aestivum*). Additionally, we evaluated two different synthetically developed wheats: a synthetic tetraploid (AADD) developed by crossing *T. monococcum* (AA) with *Aegilops tauschii* (DD), and a hexaploid synthetic (AABBDD) developed by crossing *Triticum turgidum* subsp. *dicoccoides* (AABB) and *Aegilops tauschii* (DD). Details about the genotypes are provided in Supplementary Table S1. Ten seeds of each genotype were selected, and four roll-ups were prepared for each genotype, each representing one replication. The seeds were rolled up 2.5 cm from the top edge of a sheet of brown germination paper (Anchor Paper Co., St. Paul, MN, USA). The roll-ups were placed upright in a 2.0 L beaker containing 500 ml of 0.5 mM CaSO4 solution, with each replication in a separate beaker. The beakers with the seedling roll-ups were kept in a germination chamber (30 °C in darkness at 60-70% humidity) for 48 h. After germination, the roll-ups were transferred to a growth chamber (Environmental Growth Chambers, Model GC-36, Chagrin Falls, OH44022, US) with a 16/8-h (light/dark) photoperiod (∼600 µmol m^−2^ s^−1^ photosynthetically active radiation was provided by metal halide bulbs), 40% relative humidity, and maximum/minimum temperatures of 24°C/26°C. Fifteen days post-imbibition, the plants were harvested for root phosphorus and nitrogen content, as well as for anatomy sample collection. From each replication, one healthy plant was selected from each roll-up, and two healthy seminal roots were collected from each plant. Four replications were harvested per genotype. A root segment 1.5 to 2 cm long and one cm away from the base was collected from each root, placed into a histocap (Bio Plas™ 6020, Histo Plas^TM^, California, USA), and preserved in 70% (v/v) ethanol in water. The samples were then dried using a critical point drier (Leica EM CPD 300, Leica Microsystems Inc, Buffalo Grove, IL, USA) to preserve root anatomy. Laser ablation tomography (LAT), as described by Strock et al. (2018), was used for imaging. Root anatomical phenes were measured using the ObjectJ plugin in ImageJ software. The modified Hagen-Poiseuille law (Equation 1) was applied to cross-sectional images of the segment to determine the theoretical axial hydraulic conductance (kh; kg m MPa^−1^ s^−1^), with d representing the diameter of each metaxylem vessel in meters, ρ representing the fluid density (which is equivalent to water at 20°C and has a value of 1,000 kg m^−3^), and η representing the viscosity of the fluid (which is also equal to water at 20°C and has a value of 1 × 10^−9^ MPa s^−1^) (Tyree and Ewers, 1991).

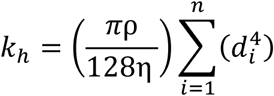

Equation 1. Modified Hagen-Poiseuille equation.

Adjacent to each plant’s anatomy samples, one 5 cm long root segment was collected for phosphorus content measurement, and another 5 cm long root segment was collected for nitrogen content measurement. For phosphorus content measurements, the segments were first dried at 64°C, then weighed to obtain dry weight. The dry root segments were ashed at 500°C for 16 hours, dissolved in 100 mM HCl, and analyzed spectrophotometrically (Murphy and Riley, 1962). Root samples collected for nitrogen content were first dried at 64°C, and root N content was measured using an elemental analyzer (Perkin Elmer 2400 Series II CHN analyzer).

### Low phosphorus and low nitrogen study

A split-plot experiment was designed with four blocks representing four replicates. Each block was divided into three sub-blocks, each representing one of the three treatments: control (optimal phosphorus and nitrogen), suboptimal phosphorus, and suboptimal nitrogen. Within each subplot, a genotype-treatment combination was randomly assigned to a plot. Four-liter pots were filled with a medium containing 10% soil, 50% sand, 20% vermiculite, and 20% perlite. For the suboptimal nitrogen treatment, soil (Typic Hapludalf, silt loam) was collected from suboptimal nitrogen fields that had not received nitrogen fertilizer for over a decade at the Russell E. Larson Agricultural Research Center in Rock Springs, Pennsylvania (77°57′W, 40°42′N). Similarly, for the suboptimal phosphorus treatment, soil (Typic Hapludalf, silt loam) was collected from suboptimal phosphorus fields that had not received phosphorus fertilizer for over two decades at the same research center. For the control treatment, soil (Typic Hapludalf, silt loam) was collected from fields that received optimum nitrogen and phosphorus. Before planting, the pots were watered to field capacity, and the seeds were planted 4 cm below the soil surface. For the first 10 days, no additional fertilization was provided. Starting from the 10th day, every other day, the control pots were given 100 ml of the optimum nutrient solution (KNO_3_ at 5000 μM, Ca(NO_3_)_2_ at 5000 μM, MgSO_4_ at 5000 μM, KH_2_PO_4_ at 2500 μM, Fe-DTPA at 150 μM, H_3_BO_3_ at 46.25 μM, MnCl_2_ at 9.14 μM, ZnSO_4_ at 0.76 μM, CuSO_4_ at 0.32 μM, and (NH_4_)_6_Mo_7_O_2_ at 0.08 μM), while suboptimal nitrogen pots were given the same solution as the control, except that Ca(NO_3_)_2_ was replaced with CaCl_2_ (5000 μM) and KCl (2220 μM), and the concentration of KNO_3_ was adjusted to 625 μM. For the suboptimal phosphorus solution, KH_2_PO_4_ was replaced with KCl (2500 μM). Throughout the experiment, the plants were grown under controlled greenhouse conditions (16 h of day at 26°C / 8 h of night at 24°C, 40%–70% relative humidity). Midday photosynthetic active radiation was ∼900–1000 μmol photons m^−2^ s^−1^. Natural light was supplemented during the day with ∼500 μmol photons m^−2^ s^−1^ from LED lamps.The plants were harvested 30 days after planting. The shoots were harvested and dried at 64°C for dry weight measurement, and the roots were washed, divided into each root class, and dried at 64°C to obtain the dry root weight.

### Evaluation of root penetration ability

#### Wax-petrolatum layer setup

For this experiment we tested two genotypes each of nine species including four diploids (*Triticum monococcum*, *Triticum urartu, Aegilops speltoides*, *Aegilops tauschii*), three tetraploids (*Triticum turgidum* subsp. *dicoccoides*, *Triticum turgidum* subsp. *dicoccon, Triticum turgidum* subsp. durum) and a hexaploid (*Triticum aestivum*). Additionally, two synthetic hexaploids (see section *Root anatomy and tissue nutrient content* for details) were generated by crossing *Triticum turgidum* subsp. *dicoccoides* and *Aegilops tauschii* were also included. Details about the genotypes are given in supplementary table S1. Wax-petrolatum layers of 2 mm thickness and strength equivalent to 1.0 MPa were utilized to mimic hard soils (Long-Xi et al., 1995). To prepare wax-petrolatum layers we mixed and melted equal amounts (w/w) of wax (Royal Oak Sales, Inc, GA30076, US) and petrolatum (Unilever, CT06611, US). The wax-petrolatum mix was poured and molded into the bottom of polyvinyl chloride cylinders (PVC, 5 cm diameter and 5 cm height) to form a 2 mm thick layer. After allowing the wax layers to harden for 4 hours, the columns were filled with growth medium mix (G1) consisting of, v/v, 50% commercial grade sand, 35% vermiculite, 15% topsoil (Hagerstown silt loam topsoil, a fine, mixed, mesic Typic Hapludalf), and 5 g/L of Osmocote plus fertilizer (Scotts-Sierra Horticultural Products Company, Marysville, Ohio, USA) consisting of (%): NO_3_^−^ (8) NH_4_^+^ (7), P (9), K (12), S (2.3), B (0.02) Cu (0.05), Fe (0.68), Mn (0.06), Mo (0.02), and Zn (0.05).

The columns were then placed on top of the 40 L containers filled with growth media (G1), with the wax-petrolatum layers in close contact with the growth medium surface. Each 40 L container represented a block and in total four containers were used in a controlled growth chamber. Control columns with no wax-petrolatum layer were also installed in each container. Control and treatment columns within a block were randomly assigned a genotype. The experimental design was a randomized complete block. One seed in each column was planted at a depth of 2 cm and the columns as well as the 40 L containers were watered to field capacity. The plants were grown in a growth chamber with a constant temperature of 24°C and approximately 600 µmol m^-2^ s^-1^ photosynthetically active radiation (PAR) was provided by metal halide lamps for four weeks. At the end of the growth period, the shoots were harvested and dried at 64°C to obtain shoot dry weight. The roots were harvested by carefully cutting the penetrated roots using scissors and washing the soil. The total number of roots, both nodal and seminal, were counted and the penetration ratio was expressed as the number of roots penetrated/total number of roots. The relative reduction in shoot dry biomass per replication was calculated as follows (shoot dry weight under control condition – shoot dry weight under hard wax-petroatum layer treatment)/ shoot dry weight under control condition).

### Hard soil penetration

To confirm the wax-petrolatum layer experiment results, we tested the penetration ability of two extreme ploidy levels i.e. diploid (*Triticum monococcum*) and hexaploid (*Triticum aestivum*). Three genotypes of each species were selected (Supplementary Table S1). Topsoil (Hagerstown silt loam, fine, mixed, mesic Typic Hapludalf) was first dried and then sieved using a 2 mm mesh sieve. Using a manual press sieved soil was compacted into columns (5 cm diameter and 10 cm height) to achieve 1.1 g cm^-3^ bulk density for the control treatment and 1.4 g cm^-3^ bulk density for the increased mechanical impedance treatment. In total four replications per genotype under each treatment were planted with each column containing one seed. The plants were grown in a growth chamber with a 16/8-h (light/dark) photoperiod (approximately 600 µmol m^-2^ s^-1^ photosynthetically active radiation (PAR) were provided by metal halide lamps), 40% relative humidity, and maximum/minimum temperatures of 24°C/26°C. Ten days post planting, columns were washed and roots from each column were scanned on a flat-bed image using an EPSON Perfection V700 PHOTO scanner, and total root length was quantified with WinRhizo software (WinRhizo Pro; Reagent Instruments Inc, Québec, Canada).

### Tip shape measurements

Plants were grown as described in the root anatomy and tissue nutrient content section. Fifteen days post planting the root tips of healthy seedlings were imaged under a stereomicroscope (Nikon SMZ1500, Nikon^®^, Tokyo, Japan)), and images were captured with a Nikon DS-Fi1 camera at 4X magnification and NIS-Elements F2.30 software (Nikon, Tokyo, Japan). The catenary curves were fit to the root shapes (Fujiwara et al., 2021) using our python language-based program. The catenary curve is described by equation 2.

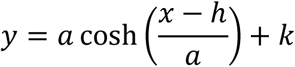

Equation 2. The catenary curve is used to describe the root tip shape. The parameter “a” defines the bluntness/roundness of a curve with an increase in “a” corresponding to increased bluntness. “h” is horizontal shift (along the x-axis) and “k” is vertical shift (along the y-axis).

### *RootSlice* simulations

In order to dissect the role of vacuolar occupancy and cortical cell size in root tissue nutrient content we used *RootSlice* (Sidhu et al., 2023)*. RootSlice* is a multicellular functional-structural model of root anatomy capable of simulating a variety of natural and artificial phenotypes facilitating the analysis, exploration, and understanding of root anatomical phenotypes. In this study we used *RootSlice* to understand the role of cortical cell size and vacuolar occupancy in regulating root tissue nutrient content. We simulated the root anatomy of three ploidy levels including diploids (*Triticum monococcumi*), one tetraploid (*Triticum turgidum subsp. dicoccon*), and a hexaploid (*Triticum aestivum*). Parameters for simulating each species are given in Supplementary Table 2 and represent the average phenotype values for each species. Cortical cell cross-sectional area for *T. monococcum*, *T. t. dicoccon,* and *T. aestivum* was set to 325, 548, and 604 µm^2^, respectively. Longitudinally, each root segment comprises of three cell layers and each cell layer was set to 100 µm long. Additionally, to highlight the effect of cortical cell size on tissue phosphorus and nutrient content, we simulated root segments of each species with all other original parameters except CCS, which was used from the hexaploid (*Triticum aestivum*). The version used to simulate each model can be found at https://github.com/iba5104/RootSlice. Input files used for this study can be found at 10.5281/zenodo.8083558. The visualization of three dimensional root models was performed using Paraview (https://vtk.org/).

### Open-SimRoot simulations

To simulate native and cultivated PPNB environments (‘cultivation’ indicating the management of desired plants through soil tillage (manual or animal-drawn), planting, weed suppression, and protection from herbivores) we used the parameters given in (Supplementary table S4, and in the following Zenodo repository 10.5281/zenodo.15271839.). Adaptation of ploidy-specific phenotypes to these environments was tested by generating three root models, a) hexaploid wheat model with actual hexaploid wheat parameters as presented in this study, tetraploid and diploid wheat models with everything kept the same as the hexaploid model except for tetraploid and diploid values of root phosphorus content, nitrogen content, root respiration, axial hydraulic conductance, and root penetration ability (Supplementary table S3). The input files required to reproduce these simulations are available in the following Zenodo repository 10.5281/zenodo.15271839.

### Statistical analysis

All data analyses presented in this study were conducted using the R programming language v 4.4.2 in RStudio (R Core Team, 2016). To test for differences in all parameters except shoot biomass reduction under low phosphorus and nitrogen (Fig. 1B) among different ploidy levels, we used analysis of variance (ANOVA) with a nested design, where Species were nested within Ploidy, and Genotypes were nested within Species, implemented using the “aov” function from the stats package in R. Following ANOVA, pairwise comparisons were performed using Tukey’s Honest Significant Difference (HSD) test via the “TukeyHSD” function from the stats package.

For testing differences in shoot biomass reduction due to low phosphorus and nitrogen (Fig. 1B), we used a non-parametric Kruskal-Wallis test, implemented using the kruskal.test() function from the stats package. To assess relationships between cortical cell size and root nitrogen, phosphorus, and respiration traits, linear models were fit using the lm() function from the stats package in R. The significance of all models including ANOVA and Kruskal-Wallis was evaluated at a significance level of α = 0.05.

## Acknowledgments

We thank Christopher Benson for providing the Poa seedlings and thank Scott Diloreto for his support with the greenhouse experiments. JSS also gratefully acknowledges Dr. Navtej Singh Bains for his insights into wheat evolution and for inspiring JSS to study wheat during his undergraduate years at Punjab Agricultural University.

## Funding

The US Department of Energy ARPA-E Award DE-AR0000821

The Howard G Buffett Foundation

The US Department of Agriculture National Institute of Food and Agriculture Hatch Appropriations Projects PEN04732, PENW-2020-03632 and Accession #:7009406.

## Author contributions

Conceptualization: JSS and JPL

Methodology: JSS, HSG, SW, MS, RJHS, SS, JPL

Investigation: JSS, SW, HSW, ILV, HR, IA Visualization: JSS

Funding acquisition: JPL and JSS Supervision: JPL

Writing – original draft: JSS

Writing – review & editing: JSS, JPL, HSG, ILV, SW, MS, RJHS, SS

## Data and materials availability

All data, code, and materials used in the analysis will be available at the zenodo repository 10.5281/zenodo.15271839.

## Supplementary Materials

Supplementary Text Fig. S1 – S3

Tables S1 – S4

