## Supplementary data for "Ploidy alters root anatomy and shapes the evolution of wheat polyploids"

**Affiliations:**

This PDF file includes:

Figures S1, S2, and S3

Tables S1, S2, S3, and S4

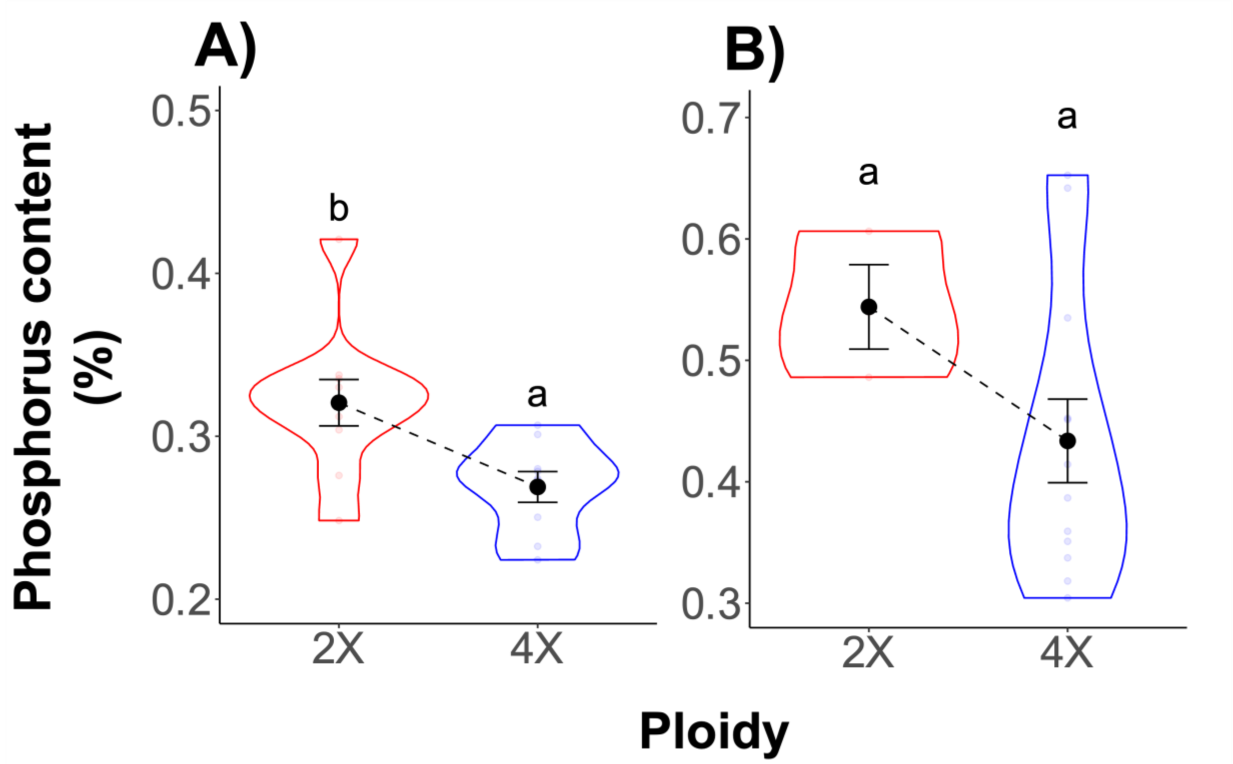


**Supplementary Figure 1. Root phosphorus content in *Poa* and *Gossypium* species.** *Poa supina* (2X, SS) is a diploid while *Poa annua* (4X, IISS) is tetraploid. *Gossypium herbaceum* (2X, AA) is diploid and *Gossypium hirsutum* (4X, AADD) is a tetraploid. The letters on top of the violin plots denote Wilcoxon rank sum exact test levels with groups having the same letters indicating no significant difference at $p \leq 0.05$, while those with different letters indicate a significant difference.

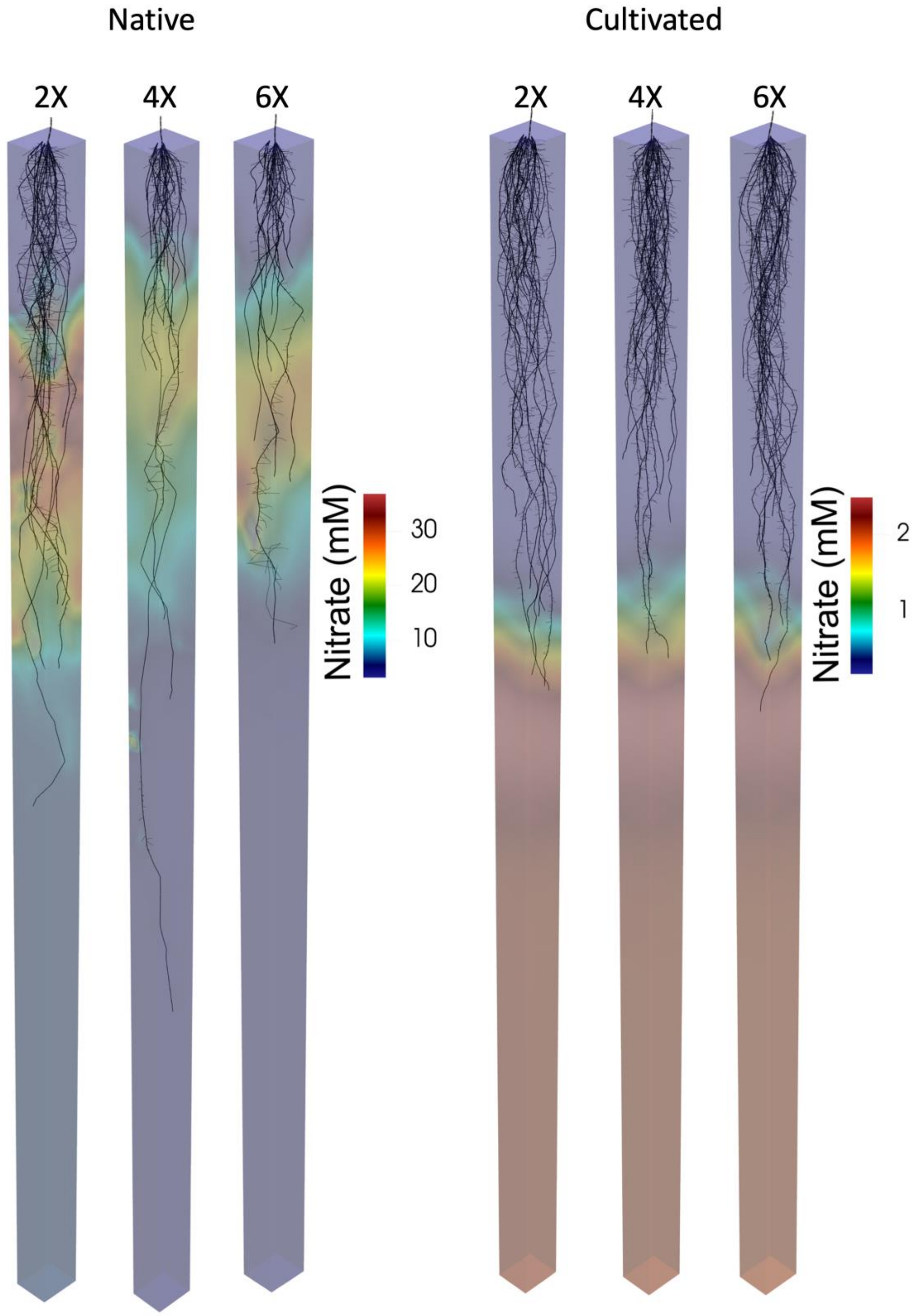
Native
Cultivated
2X
4X
6X
2X
4X
6X
Nitrate (mM)
30
20
10
Nitrate (mM)
2
1

**Supplementary Figure S2.** *Simulated root growth of wheat with different ploidy levels in Pre-Pottery Neolithic native (~10,000 BP) and cultivated (~8,000 BP) environments.* Root architecture and soil conditions were simulated using the functional–structural model *OpenSimRoot v2*. Nitrate in the soil solution, as influenced by mineralization, diffusion, mass flow, and root uptake is visualized as a color gradient, expressed in mmol $L^{-1}$ of soil solution. In native (i.e. uncultivated) soil environments, diploid phenotypes exhibit better root growth compared to tetraploid and hexaploid phenotypes. In contrast, in cultivated environments, tetraploid and hexaploid phenotypes show superior root development.

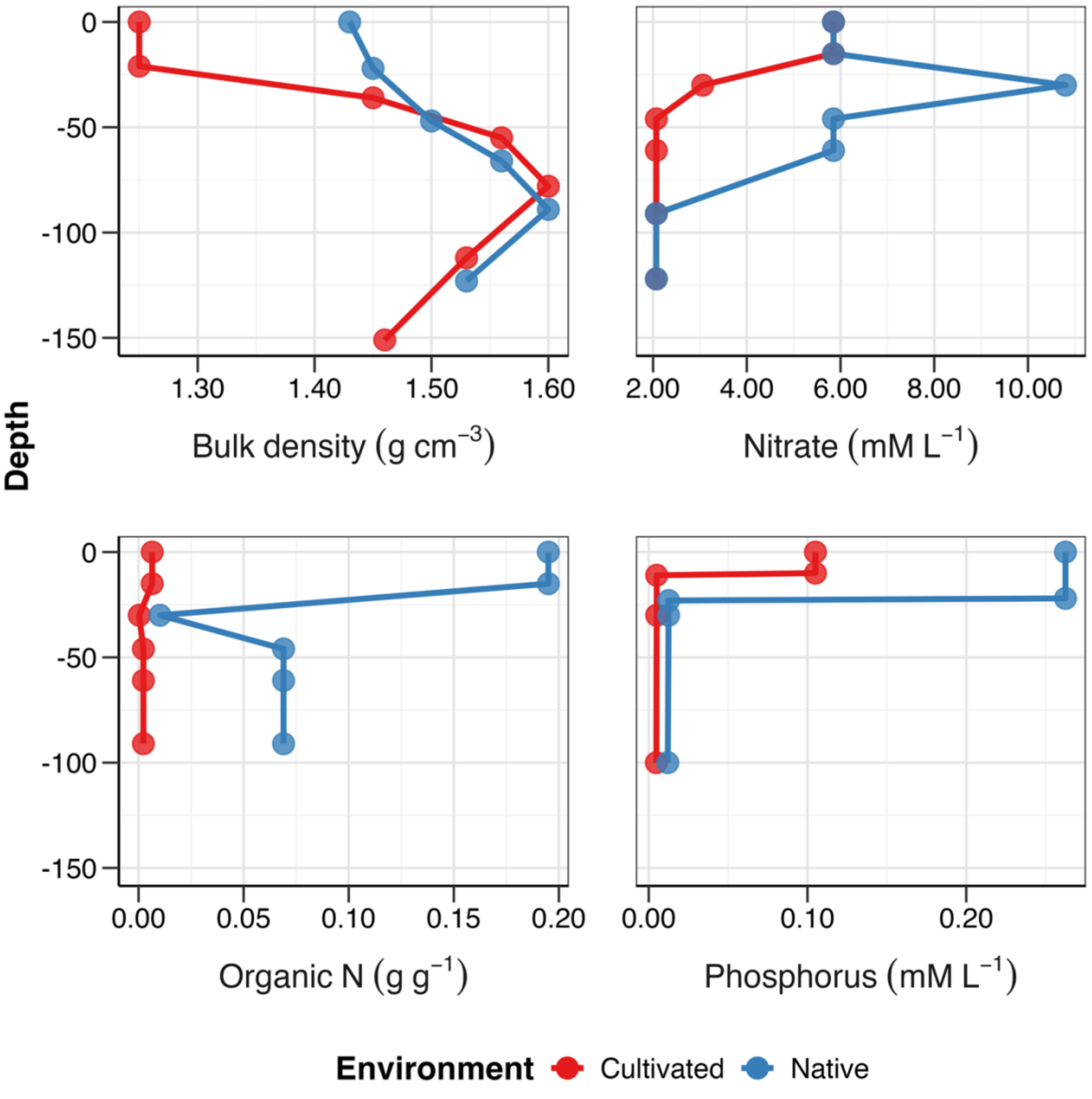


**Supplementary Figure S3.** Soil profiles for the neolithic native (i.e., uncultivated) and cultivated environments as parameterized in *OpenSimRoot v2*.

**Table S1. Details about the genotypes used for different experiments in this study.**

| GENOTYPE | SPECIES | EXPERIMENT |
|---|---|---|
| **PI 173614** | *Aegilops speltoides* | Respiration, Root Penetration Ability |
| **PI 369618** | *Aegilops speltoides* | Root Penetration Ability |
| **PI 560747** | *Aegilops speltoides* | Respiration |
| **AE 525** | *Aegilops tauschii* | Respiration, Tip Shape, Synthetics Physiology |
| **AT-7** | *Aegilops tauschii* | Anatomy, Synthetics Physiology |
| **AT-8** | *Aegilops tauschii* | Anatomy, Synthetics Physiology |
| **CLAE 8** | *Aegilops tauschii* | Nutrient Sensitivity, Synthetics Physiology |
| **TA 1644** | *Aegilops tauschii* | Anatomy, Synthetics Physiology |
| **TA 1645** | *Aegilops tauschii* | Respiration, Synthetics Physiology |
| **TA 1669** | *Aegilops tauschii* | Nutrient Sensitivity, Root Penetration Ability, Synthetics Physiology |
| **TA 2549** | *Aegilops tauschii* | Root Penetration Ability |
| **TA1644** | *Aegilops tauschii* | Tip Shape |
| **TA1645** | *Aegilops tauschii* | Tip Shape |
| **SYN86** | *SYN_AABBDD* | Tip Shape, Synthetics Physiology |
| **SYN89** | *SYN_AABBDD* | Synthetics Physiology |
| **TA 3361** | *SYN_AADD* | Synthetics Physiology |
| **TA3361** | *SYN_AADD* | Tip Shape |
| **BERKUT** | *Triticum aestivum* | Anatomy, Nutrient Sensitivity, Respiration, Tip Shape |
| **BRIGGS** | *Triticum aestivum* | Anatomy, Nutrient Sensitivity, Tip Shape |
| **CHINESE SPRING** | *Triticum aestivum* | Tip Shape |
| **CHINESE SPRING** | *Triticum aestivum* | Root Penetration Ability, Compacted Soil Experiment |
| **OVERLAND** | *Triticum aestivum* | Root Penetration Ability, Compacted Soil Experiment |
| **PREVAIL** | *Triticum aestivum* | Anatomy, Respiration |
| **RB07** | *Triticum aestivum* | Anatomy, Respiration, Root Penetration Ability, Compacted Soil Experiment, Tip Shape |
| **SONORA** | *Triticum aestivum* | Root Penetration Ability |
| **PI 167526** | *Triticum monococcum* | Root Penetration Ability |
| **PI 167589** | *Triticum monococcum* | Nutrient Sensitivity |
| **PI 167625** | *Triticum monococcum* | Compacted Soil Experiment |
| **PI 221413** | *Triticum monococcum* | Tip Shape |
| **PI 266844** | *Triticum monococcum* | Nutrient Sensitivity |
| **PI 427798** | *Triticum monococcum* | Synthetics Physiology |
| **PI427734** | *Triticum monococcum* | Synthetics Physiology |
| **TA 1988** | *Triticum monococcum* | Anatomy, Respiration, Synthetics Physiology |
| **TA 2031** | *Triticum monococcum* | Anatomy, Compacted Soil Experiment, Synthetics Physiology |
| **TA 2708** | *Triticum monococcum* | Compacted Soil Experiment |
| **TA 4342** | *Triticum monococcum* | Synthetics Physiology |
| **TA 4342-L95** | *Triticum monococcum* | Anatomy |

| | | |
|---|---|---|
| **TA 4342-L96** | *Triticum monococcum* | Anatomy |
| **TA 4343 - L95** | *Triticum monococcum* | Synthetics Physiology |
| **TA 4343-L95** | *Triticum monococcum* | Respiration |
| **TA2027** | *Triticum monococcum* | Tip Shape |
| **TA2708** | *Triticum monococcum* | Tip Shape |
| **TA4342-L95/L96** | *Triticum monococcum* | Root Penetration Ability |
| **G-148** | *Triticum turgidum subsp. dicoccoides* | Tip Shape |
| **G148** | *Triticum turgidum sub sp. dicoccoides* | Respiration, Synthetics Physiology |
| **G41M** | *Triticum turgidum sub sp. dicoccoides* | Respiration, Synthetics Physiology |
| **PI 479780** | *Triticum turgidum subsp. dicoccoides* | Root Penetration Ability |
| **PI 479781** | *Triticum turgidum subsp. dicoccoides* | Root Penetration Ability |
| **PI 538636** | *Triticum turgidum subsp. dicoccoides* | Tip Shape |
| **PI 538637** | *Triticum turgidum sub sp. dicoccoides* | Respiration |
| **PI 538640** | *Triticum turgidum subsp. dicoccoides* | Tip Shape |
| **PI 656868** | *Triticum turgidum sub sp. dicoccoides* | Respiration |
| **CLTR 4013** | *Triticum turgidum subsp. dicoccon* | Respiration, Root Penetration Ability |
| **CLTR 4013** | *Triticum turgidum subsp. dicoccon* | Anatomy |
| **E-TDCN-5** | *Triticum turgidum subsp. dicoccon* | Anatomy |
| **KU-124** | *Triticum turgidum subsp. dicoccon* | Tip Shape |
| **MGS 29311** | *Triticum turgidum subsp. dicoccon* | Anatomy |
| **MGS293/1** | *Triticum turgidum subsp. dicoccon* | Respiration |
| **PI 355497** | *Triticum turgidum subsp. dicoccon* | Root Penetration Ability |
| **PI 94625** | *Triticum turgidum subsp. dicoccon* | Tip Shape |
| **PI 134951** | *Triticum turgidum subsp. durum* | Root Penetration Ability |
| **PI 156263** | *Triticum turgidum subsp. durum* | Anatomy |
| **PI 185233** | *Triticum turgidum subsp. durum* | Respiration |
| **PI 192003** | *Triticum turgidum subsp. durum* | Anatomy, Tip Shape |

| | | |
|---|---|---|
| **PI 192843** | *Triticum turgidum subsp. durum* | Respiration |
| **PI 324481** | *Triticum turgidum subsp. durum* | Root Penetration Ability, Tip Shape |
| **PI 388035** | *Triticum turgidum subsp. durum* | Anatomy |
| **PI 60741** | *Triticum turgidum subsp. durum* | Anatomy |
| **PI 662222** | *Triticum urartu* | Respiration, Root Penetration Ability |
| **PI 662229** | *Triticum urartu* | Root Penetration Ability |
| **PI 662244** | *Triticum urartu* | Respiration |
| **PI 662251** | *Triticum urartu* | Respiration |

**Table S2. *RootSlice* parameterization to simulate root anatomical phenotypes of different species of wheat and its ancestors.**

| *Species* | *Ploidy* | *Cortical cell cross sectional area (CCS) ($um^2$)* | *Cortical cell file number* | *Cell length (um)* | *Stele diameter (um)* |
|---|---|---|---|---|---|
| *T. monococcum* | 2X | 325 | 6 | 100 | 280 |
| *T. t. dicoccon* | 4X | 548 | 6 | 100 | 322 |
| *T. aestivum* | 6X | 604 | 6 | 100 | 342 |

**Table S3. *OpenSimRoot v2* parameterization to simulate root phenotypes for each ploidy level.**

| **Root parameter** | ***6X - T. aestivum*** | ***4X T. t. dicoccon*** | ***2X T. monococcum*** |
|---|---|---|---|
| Tissue Density (g $cm^{-3}$) dry weight | Mean = 0.074, SE = 0.019 | Mean = 0.078, SE = 0.027 | Mean = 0.110, SE = 0.051 |
| Root respiration (nmol CO2 per g per sec) | Mean = 3.32, SE = 0.227 | Mean = 3.68, SE = 0.125 | Mean = 5.98, SE = 0.067 |
| Root P content (%) | Mean = 0.89, SE = 0.043 | Mean = 1.13, SE = 0.043 | Mean = 1.19, SE = 0.064 |
| Root N content (%) | Mean = 1.19, SE = 0.099 | Mean = 1.42, SE = 0.10 | Mean = 1.66, SE = 0.11 |
| Root penetration ability (Penetration ratio) | Mean = 0.19, SE = 0.05 | Mean = 0.351, SE = 0.056 | Mean = 0.52, SE = 0.10 |

**Table S4. *OpenSimRoot v2* soil parameterization summary used to simulate Neolithic cultivated and native environments.**

| *Environment* | *Native* | *Cultivated* |
|---|---|---|
| *Phosphorus* | High | Low |
| *Nitrogen* | High | Low |
| *Water* | High | Low |
| *Penetration resistance* | High | Low |
| *Atmospheric $CO_2$ ppm* | 270 PPM | 270 PPM |
